# Characterization of accessory genes in coronavirus genomes

**DOI:** 10.1101/2020.05.26.118208

**Authors:** Christian J. Michel, Claudine Mayer, Olivier Poch, Julie D. Thompson

## Abstract

The Covid19 infection is caused by the SARS-CoV-2 virus, a novel member of the coronavirus (CoV) family. CoV genomes code for a ORF1a / ORF1ab polyprotein and four structural proteins widely studied as major drug targets. The genomes also contain a variable number of open reading frames (ORFs) coding for accessory proteins that are not essential for virus replication, but appear to have a role in pathogenesis. The accessory proteins have been less well characterized and are difficult to predict by classical bioinformatics methods. We propose a computational tool GOFIX to characterize potential ORFs in virus genomes. In particular, ORF coding potential is estimated by searching for enrichment in motifs of the *X* circular code, that is known to be over-represented in the reading frames of viral genes. We applied GOFIX to study the SARS-CoV-2 and related genomes including SARS-CoV and SARS-like viruses from bat, civet and pangolin hosts, focusing on the accessory proteins. Our analysis provides evidence supporting the presence of overlapping ORFs 7b, 9b and 9c in all the genomes and thus helps to resolve some differences in current genome annotations. In contrast, we predict that ORF3b is not functional in all genomes. Novel putative ORFs were also predicted, including a truncated form of the ORF10 previously identified in SARS-CoV-2 and a little known ORF overlapping the Spike protein in Civet-CoV and SARS-CoV. Our findings contribute to characterizing sequence properties of accessory genes of SARS coronaviruses, and especially the newly acquired genes making use of overlapping reading frames.

## Introduction

Coronaviruses (CoVs) cause respiratory and intestinal infections in animals and humans (Cui et al., 2019). They were not considered to be highly pathogenic to humans until the last two decades, which have seen three outbreaks of highly transmissible and pathogenic coronaviruses, including SARS-CoV (severe acute respiratory syndrome coronavirus), MERS-CoV (Middle East respiratory syndrome coronavirus), and SARS-CoV-2 (which causes the disease COVID-19). Other human coronaviruses (such as HCoV-NL63, HCoV-229E, HCoV-OC43 or HKU1) generally induce only mild upper respiratory diseases in immunocompetent hosts, although some may cause severe infections in infants, young children and elderly individuals (Cui et al., 2019).

Extensive studies of human coronaviruses have led to a better understanding of coronavirus biology. Coronaviruses belong to the family *Coronaviridae* in the order nidovirales. Whereas MERS-CoV is a member of the *Merbecovirus* subgenus, phylogenetic analyses indicated that SARS-CoV-2 clusters with SARS-CoV in the *Sarbecovirus* subgenus (Coronaviridae Study Group of the International Committee on Taxonomy of Viruses, 2020). All human coronaviruses are considered to have animal origins. SARS-CoV, MERS-CoV and SARS-CoV-2 are assumed to have originated in bats (Cui et al., 2019). It is widely believed that SARS-CoV and SARS-CoV-2 were transmitted directly to humans from market civets and pangolin, respectively, based on the sequence analyses of CoV isolated from these animals and from infected patients.

All members of the coronavirus family are enveloped viruses that possess long positive-sense, single-stranded RNA genomes ranging in size from 27–33 kb. The coronavirus genomes encode five major open reading frames (ORFs), including a 5’ frameshifted polyprotein (ORF1a/ORF1ab) and four canonical 3’ structural proteins, namely the spike (S), envelope (E), membrane (M) and nucleocapsid (N) proteins, which are common to all coronaviruses (Ashour et al., 2020). In addition, a number of subgroup-specific accessory genes are found interspersed among, or even overlapping, the structural genes. Overlapping genes originate by a mechanism of overprinting, in which nucleotide substitutions in a pre-existing frame induce the expression of a novel protein in an alternative frame. The accessory proteins in coronaviruses vary in number, location and size in the different viral subgroups, and are thought to contain additional functions that are often not required for virus replication, but are involved in pathogenicity in the natural host (Schaecher and Pekosz, 2009; Liu et al., 2014).

In the face of the ongoing COVID-19 pandemic, extensive worldwide research efforts have focused on identifying coronavirus genetic variation and selection (e.g. Cagliani et al., 2020; Khailany et al., 2020; Wang et al., 2020), in order to understand the emergence of host/tissue specificities and to help develop efficient prevention and treatment strategies. These studies are complemented by structural genomics (e.g. Jin et al., 2020; Hussain et al., 2020; Srinivasan et al., 2020), as well as transcriptomics (Kim et al., 2020) and interactomics studies (Gordon et al., 2020) of the structural and putative accessory proteins.

However, there have been less studies of accessory proteins, for two main reasons (Yuen et al., 2020). First, accessory proteins are often not essential for viral replication or structure, but play a role in viral pathogenicity or spread by modulating the host interferon signaling pathways for example. This has led to some contradictory experimental results concerning the presence or functionality of accessory proteins. For instance, in a recent experiment (Gordon et al., 2020) to characterize SARS-CoV-2 gene functions, 9 predicted accessory protein ORFs (3a, 3b, 6, 7a, 7b, 8, 9b, 9c, 10) were codon optimized and successfully expressed in human cells, with the exception of ORF3b. However, another recent study using DNA nanoball sequencing (Kim et al., 2020) concluded that the SARS-CoV-2 expresses only five canonical accessory ORFs (3a, 6, 7a, 7b, 8).

Second, bioinformatics approaches for the prediction of accessory proteins are challenged by their complex nature as short, overlapping ORFs. Such proteins are known to have biased amino acid sequences compared to non-overlapping proteins (Pavesi et al., 2018). In addition, the homology-based approaches widely used to predict ORFs in genomes are less useful here, because many accessory proteins are lineage-or subgroup-specific. Thus, many state of the art viral genome annotation systems, such as Vgas (Zhang et al., 2019), only predict overlapping proteins if homology information is available. Other methods have been developed dedicated specifically to the *ab initio* prediction of overlapping genes, for example based on multiple sequence alignments and statistical estimates of the degree of variability at synonymous sites (Firth, 2014) or sequence simulations and calculation of expected ORF lengths (Schlub et al., 2018).

Here, we propose a computational tool GOFIX (Gene prediction by Open reading Frame Identification using *X* motifs) to predict potential ORFs in virus genomes. Using a complete viral genome as input, GOFIX first locates all potential ORFs, defined as a region delineated by start and stop codons. In order to predict functional ORFs, GOFIX calculates the enrichment of the ORFs in *X* motifs, i.e. motifs of the *X* circular code (Arquès and Michel, 1996), a set of 20 codons that are over-represented in the reading frames of genes from a wide range of organisms. For example, in a study of 299,401 genes from 5217 viruses (Michel, 2017), including double stranded and single stranded DNA and RNA viruses, codons of the *X* circular code were found to occur preferentially in the reading frame of the genes. This is an important property of viral genes, since it has been suggested that *X* motifs at different locations in a gene may assist the ribosome to maintain and synchronize the reading frame (Dila et al., 2019). An initial evaluation test of the GOFIX method on a large set of 80 virus genomes (Pavesi et al., 2018) showed that it achieves high sensitivity and specificity for the prediction of experimentally verified overlapping proteins (manuscript in preparation). A major advantage of our approach is that it requires only the sequence of the studied genome and does not rely on any homology information. This allows us to detect novel ORFs that are specific to a given lineage.

We applied GOFIX to study the SARS-CoV-2 genome and related SARS genomes, with a main focus on the accessory proteins. Using the extensive experimental data concerning the SARS-CoV genome and the expressed ORFs, we first show that the reading frames of the SARS-CoV ORFs are enriched in *X* motifs, including most of the overlapping accessory proteins. Exceptions include ORF3b and ORF8b which may not be functional. Then, we use GOFIX to predict and compare putative genes in related genomes of SARS-like viruses from bat, civet and pangolin hosts as well as human SARS-CoV-2.

## Materials and methods

### Genome sequences

Viral genome sequences were downloaded from the Genbank database, as shown in Table 1. The Genbank reference genomes were used as representative genomes for SARS-CoV and SARS-CoV-2. For the Bat-CoV, Cive-Cov and Pangolin-Cov genomes, we selected well annotated Genbank entries having the highest number of annotated ORFs. All CDS annotations were extracted from the Genbank files, and ORF names were standardized according to the SARS-CoV-2 nomenclature (Table 2).

**Table 1.** Genome sequences selected for the current study. Note that the SARS-CoV strain hTor02 is from humans infected during the middle and late phases of the SARS epidemic of 2013, and has a deletion of 29 nucleotides in the region of ORF8.

|  | Description | Genbank accession number |
| --- | --- | --- |
| Bat-CoV | Bat SARS-like coronavirus isolate As6526 | KY417142 |
| Civet-CoV | Civet SARS coronavirus civet007 | AY572034 |
| SARS-CoV | Human severe acute respiratory syndrome-related coronavirus strain hTor02 | NC_004718 |
| Pangolin-CoV | Pangolin coronavirus isolate PCoV_GX-P2V | MT072864 |
| SARS-CoV-2 | Human severe acute respiratory syndrome coronavirus 2 isolate Wuhan-Hu-1 | MT072688 |

**Table 2.** CDS annotations extracted from Genbank, with ORF names standardized according to the SARS-CoV-2 nomenclature. * For convenience, ORF1ab is split into 2 regions corresponding the ORF1ab gene regions upstream and downstream of the frameshift. ** SARS-CoV annotation for ORF9c was propagated from Genbank entry AY274119: SARS-CoV isolate Tor2, where it is annotated as ORF14. *** SARS-CoV-2 annotations for ORF3b, ORF7b, ORF9b and ORF9c were propagated from Genbank entry MN985325: Severe acute respiratory syndrome coronavirus 2 isolate 2019-nCoV/USA-WA1/2020.

| Bat-CoV |  |  |  | Civet-CoV |  |  |  | SARS-CoV |  |  |  |
| --- | --- | --- | --- | --- | --- | --- | --- | --- | --- | --- | --- |
| Name | Start | Stop | Length | Name | Start | Stop | Length | Name | Start | Stop | Length |
| ORF1a* | 265 | 13398 | 13134 | ORF1a | 239 | 13366 | 13128 | ORF1a | 265 | 13398 | 13134 |
| ORF1b* | 13398 | 21485 | 8086 | ORF1b | 13366 | 21459 | 8092 | ORF1b | 13398 | 21485 | 8086 |
| S | 21492 | 25217 | 3726 | S | 21466 | 25233 | 3768 | S | 21492 | 25259 | 3768 |
| ORF3a | 25227 | 26051 | 825 | ORF3a | 25242 | 26066 | 825 | ORF3a | 25268 | 26092 | 825 |
| ORF3b | 25648 | 25992 | 345 | ORF3b | 25663 | 26127 | 465 | ORF3b | 25689 | 26153 | 465 |
| E | 26076 | 26306 | 231 | E | 26091 | 26321 | 231 | E | 26117 | 26347 | 231 |
| M | 26357 | 27022 | 666 | M | 26372 | 27037 | 666 | M | 26398 | 27063 | 666 |
| ORF6 | 27033 | 27224 | 192 | ORF6 | 27048 | 27239 | 192 | ORF6 | 27074 | 27265 | 192 |
| ORF7a | 27232 | 27600 | 369 | ORF7a | 27247 | 27615 | 369 | ORF7a | 27273 | 27641 | 369 |
| ORF7b | 27597 | 27731 | 135 | ORF7b | 27612 | 27746 | 135 | ORF7b | 27638 | 27772 | 135 |
| ORF8 | 27738 | 28103 | 366 | ORF8 | 27753 | 28121 | 369 | ORF8a | 27779 | 27898 | 120 |
|  |  |  |  |  |  |  |  | ORF8b | 27864 | 28118 | 255 |
| N | 28118 | 29386 | 1269 | N | 28123 | 29391 | 1269 | N | 28120 | 29388 | 1269 |
|  |  |  |  | ORF9b | 28133 | 28429 | 297 | ORF9b | 28130 | 28426 | 297 |
|  |  |  |  | ORF9c | 28586 | 28798 | 213 | ORF9c** | 28583 | 28793 | 211 |

Table 2. CDS annotations extracted from Genbank, with ORF names standardized according to the SARS-CoV-2 nomenclature.
| Pangolin-CoV |  |  |  | SARS-CoV-2 |  |  |  |
| --- | --- | --- | --- | --- | --- | --- | --- |
| Name | Start | Stop | Length | Name | Start | Stop | Length |
| ORF1a | 249 | 13427 | 13179 | ORF1a | 251 | 13453 | 13203 |
| ORF1b | 13427 | 21514 | 8086 | ORF1b | 13453 | 21538 | 8086 |
| S | 21522 | 25331 | 3810 | S | 21521 | 25369 | 3849 |
| ORF3a | 25341 | 26168 | 828 | ORF3a | 25378 | 26205 | 828 |
|  |  |  |  | ORF3b*** | 25509 | 25680 | 172 |
| E | 26193 | 26420 | 228 | E | 26230 | 26457 | 228 |
| M | 26468 | 27136 | 669 | M | 26508 | 27176 | 669 |
| 6 | 27147 | 27332 | 186 | ORF6 | 27187 | 27372 | 186 |
| 7a | 27339 | 27704 | 366 | ORF7a | 27379 | 27744 | 366 |
| 7b | 27701 | 27832 | 132 | ORF7b*** | 27741 | 27872 | 130 |
| 8 | 27839 | 28202 | 366 | ORF8 | 27879 | 28244 | 366 |
| N | 28218 | 29471 | 1254 | ORFN | 28259 | 29518 | 1260 |
|  |  |  |  | ORF9b*** | 28269 | 28562 | 294 |
|  |  |  |  | ORF9c*** | 28719 | 28940 | 222 |
|  |  |  |  | ORF10 | 29543 | 29659 | 117 |

### Definition of X motif enrichment (XME) scores

The *X* circular code contains the following 20 codons

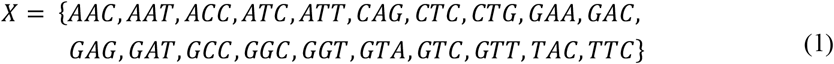

and has several strong mathematical properties (Arquès and Michel, 1996). In particular, it is self-complementary, i.e. 10 trinucleotides of *X* are complementary to the other 10 trinucleotides of *X*, and it is a circular code. A circular code is defined as a set of words such that any motif obtained from this set, allows to retrieve, maintain and synchronize the reading frame.

An *X* motif *m* is defined as a word containing only codons from the *X* circular code (1) with length |*m*| ≥ 3 codons and cardinality (i.e. number of unique codons) *c* ≥ 2 codons. The minimal length |*m*| = 3 codons was chosen based on a previous study showing that the probability of retrieving the reading frame with an *X* motif of at least 3 codons is 99.9% (Michel, 2012). The class of *X* motifs with cardinality c < 2 are excluded here because they are mostly associated with the “pure” trinucleotide repeats often found in non-coding regions of genomes (El Soufi and Michel, 2017).

The total length *XL_f_* of all *X* motifs *m_f_* of nucleotide length |*m_f_*| in a frame *f* (the reading frame or one of the 2 shifted frames) of a nucleotide sequence *s* is defined as:

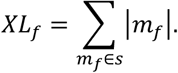

Then the *X* motif enrichment *XME_f_* in a frame *f* of a sequence *s* of nucleotide length *l* is defined as:

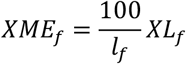

where for non-overlapping ORFs: *l_f_* = *l*, and for overlapping ORFs: *l_f_* = *l* — *XL_g_* where *XL_g_* is the total length of all *X* motifs in the overlapped frame *g*.

Finally, for an ORF of length *l* and associated with a reading frame *f* the *X* motif enrichment score *XME* is defined as:

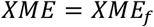

### GOFIX method

The GOFIX method will be described in detail in a separate manuscript. Briefly, the method consists of two main steps:

Identification of all potential ORFs. Using the complete genome sequences as input, all potential ORFs in the positive sense are located, defined as a sequence region starting with a start codon (AUG) and ending with a stop codon (UAA, UAG, UGA). For a given region, if alternative start codons are found, the longest ORF is selected. In this study, we selected all ORFs having a minimum length of 120 nucleotides (40 amino acids).
Calculation of *X* motif enrichment scores. For each potential ORF, all *X* motifs in the nucleotide sequence are identified in the three positive sense frames *f* using the computational method described in El Soufi and Michel (2016). For each identified potential ORF, the *X* motif enrichment (XME_f_ and XME) scores are calculated as defined above. Based on our benchmark studies (data not shown) of experimentally validated ORFs in a large set of 80 genomes (Pavesi et al., 2018), we set the threshold for prediction of a functional ORF to be *XME* ≥ 5.

## Results

### Initial study of SARS-CoV reference genome

We first analyzed the complete genome of the well-studied SARS-CoV and plotted the *X* motif enrichment (XME_f_) scores calculated in a sliding window of 150 nucleotides for each of the three positive sense frames (Fig. 1). We then mapped the ORF1ab, the four structural proteins (S, E, M, N), and the nine generally accepted accessory genes (3a, 3b, 6, 7a, 7b, 8a, 8b, 9b, 9c) to the *X* enrichment plot.

**Fig. 1.**
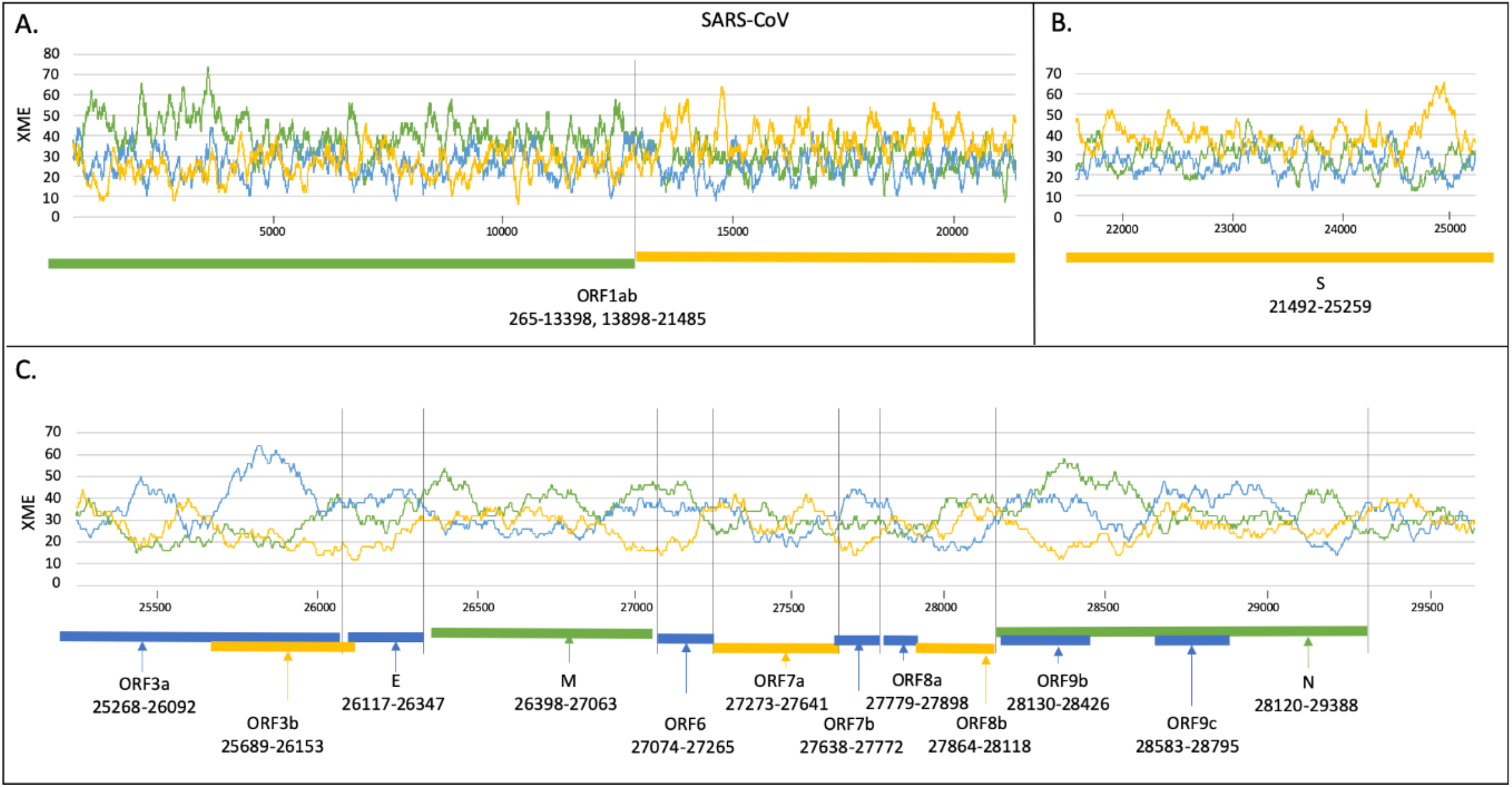
*X* motif enrichment (XME_f_) scores in the three frames *f* = 0, 1 and 2 (green, blue, yellow respectively) of the SARS-CoV genome, using a sliding window of length 150 nucleotides. Genomic organization of known ORFs is shown underneath the plots. **A.** Polyprotein gene ORF1ab. **B.** Spike protein. **C.** C-terminal structural and accessory proteins. The colors used in the enrichment plot and in the boxes representing ORFs (green, blue, yellow) indicate the three frames 0,1 and 2 respectively.

We observe a tendency for the reading frames of the SARS-CoV ORFs to be enriched in *X* motifs. For example, ORF1ab is the longest ORF, encoding a polyprotein, which is translated by a −1 programmed ribosomal frameshift at position 13398. Sequences upstream and downstream of the frameshift are enriched in *X* motifs in the corresponding reading frame (green and yellow plots respectively in Fig. 1A). Other ORFs enriched in *X* motifs in the reading frame include the S protein (yellow plot in Fig. 1B) and the E and M proteins (blue and green plots respectively in Fig. 1C). The S, E and M ORFs are conserved in all coronavirus genomes and code for structural proteins that together create the viral envelope.

The case of overlapping ORFs is more complex. For example, the last structural protein coded by the N ORF is overlapped by two accessory genes: ORF9b and ORF9c. The sequence regions containing the overlapping ORFs are characterized by an enrichment in *X* motifs in the 2 frames (green and blue plots in Fig. 1C).

### Characterization of known accessory genes in SARS-CoV

The SARS-CoV genome is known to contain four structural proteins and nine accessory proteins, namely ORFs 3a, 3b, 6, 7a, 7b, 8a, 8b, 9b and 9c. To verify that our approach can predict the accessory genes in coronavirus genomes, we used GOFIX to identify all potential ORFs in the complete SARS-CoV genome and calculate their *X* enrichment. Fig. 2 shows the *X* motif enrichment (XME_f_) scores calculated by GOFIX for the identified ORFs in the 3’ terminal region of the SARS-CoV genome.

**Fig. 2.**
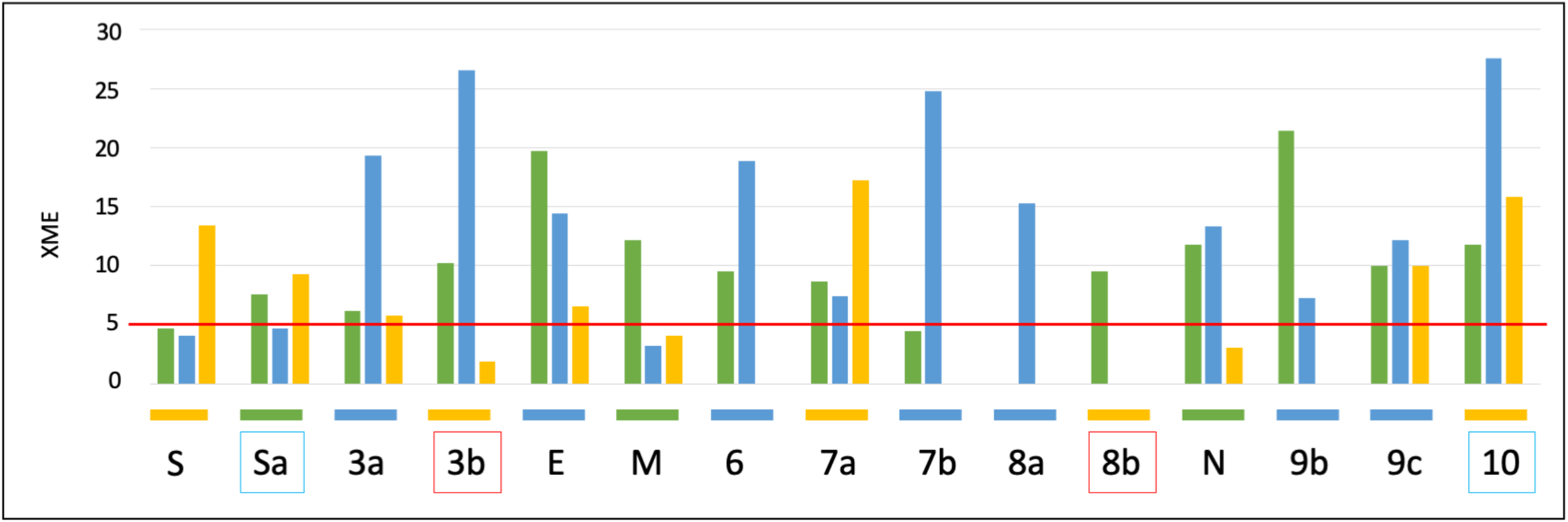
XME_f_ scores calculated by GOFIX for potential ORFs in the 3’ terminal region of the SARS-CoV genome, in the three frames *f* = 0, 1 and 2 (green, blue, yellow respectively). For clarity, only Genbank annotated ORFs or new ORFs predicted in this work are shown. The red line represents the threshold value XME=XME_f_=5 (where *f* is the reading frame) for the prediction of a functional ORF. Known ORFs are indicated below the histogram using the color corresponding to the ORF reading frame. Known ORFs not predicted to be functional by GOFIX are outlined in red. Novel ORFs predicted by GOFIX are outlined in blue.

The overall performance of GOFIX is shown in Table 3. Initially, GOFIX found 25 potential ORFs (delineated by start and stop codons) in the 3’ region (21492-29751) of SARS-CoV. Twelve of these 25 potential ORFs were predicted to be non-functional (see Methods), including 10 unknown ORFs mostly overlapping the S protein. Two previously annotated ORFs were also predicted to be non-functional, namely ORF3b (XME=1.9) and ORF8b (XME=0.0) that are discussed in detail below.

**Table 3.** Prediction performance of the GOFIX method on the set of known ORFs in the SARS-CoV genome.

|  | Predicted:<br>YES | Predicted:<br>NO | Total |
| --- | --- | --- | --- |
| Known ORF | 11 | 2 | 13 |
| Unknown ORF | 2 | 10 | 12 |
| Total | 13 | 12 | 25 |
|  | Sensitivity=0.85 | Specificity=0.83 |  |

GOFIX predicts that 13 of the 25 potential ORFs are functional (with XME>5). These include 11 previously annotated ORFs, namely S, 3a, E, M, 6, 7a, 7b, 8a, N, 9b, 9c. Two novel ORFs are also predicted by the GOFIX method: ORF10 (XME=15.8) is located downstream of the N gene (29415-29496) and a new ORF we called ORFSa (XME=7.6) that overlaps the S gene (22732-22928). These novel ORFs are discussed in more detail below.

### Comparative analyses of accessory proteins in coronavirus genomes

Having evaluated the GOFIX method on the well-studied SARS-CoV genome, we then used it to characterize and compare the accessory proteins in representative strains of five coronavirus genera, including SARS-CoV, SARS-CoV-2 and three viruses from animal hosts with SARS-CoV-like infections. Bat is considered to be the most likely host origin of SARS-CoV and SARS-CoV-2. It is generally considered that transmission to humans occurred *via* an intermediate host. For SARS-CoV, civets probably acted as the intermediate host, while pangolin has been proposed as the intermediate host in SARS-CoV-2 animal-to-human transmission (Zhang et al., 2020). For each of the five genomes, we used GOFIX to predict all potential ORFs in the complete genomes and calculated the *X* motif enrichment (XME) scores for each ORF. Fig. 3 gives an overview of the predicted ORFs in each genome, confirming for example that the structural proteins S, E, M and N, as well as the accessory proteins ORF6, ORF7a and ORF7b are conserved and have XME scores above the defined threshold XME=5. However, important differences in XME scores are observed for the remaining accessory protein ORFs.

**Fig 3.**
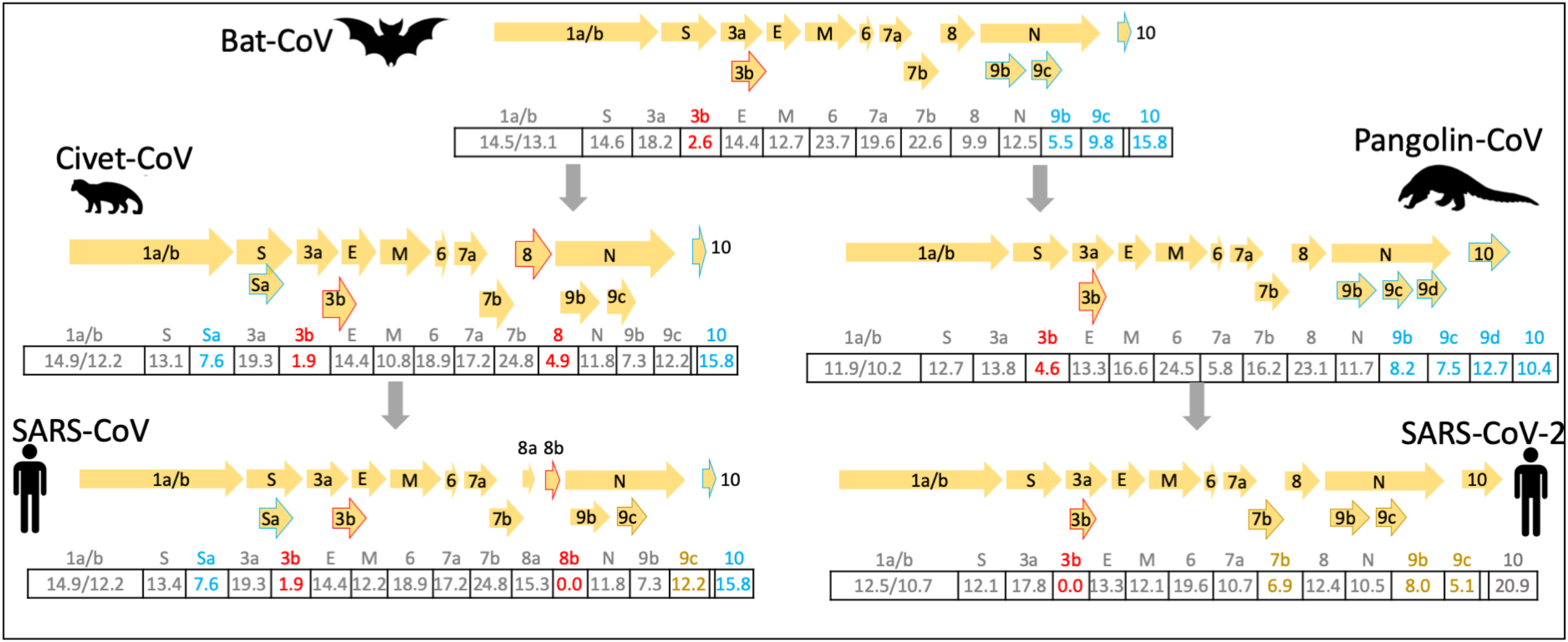
Prediction of ORFs in representative SARS-like coronavirus genomes. A schema is provided for each genome, showing the Genbank annotated ORFs and new ORFs predicted in this work. The numbers in the tables below each schema indicate the XME scores of each ORF. Genbank annotated ORFs that are not predicted to be functional by the GOFIX method are highlighted in red. Novel ORFs predicted by GOFIX are shown in blue. ORFs with conflicting annotations in Genbank, but predicted by GOFIX are shown in brown. Note that ORF3b in Civet-CoV and SARS-CoV is not homologous to ORF3b in Pangolin-CoV and SARS-CoV-2.

### ORF3b may not code for a functional protein in all CoVs

ORF3a codes for the largest accessory protein that comprises 274-275 amino acids (Fig. 4). In SARS-CoV, ORF3a is not required for virus replication, but contributes to pathogenesis by mediating trafficking of Spike (S protein) (Schaecher and Pekosz, 2009). It is efficiently expressed on the cell surface, and was easily detected in a majority of SARS patients. The XME scores for ORF3a in all the genomes range from 13.8-19.3, *i.e.* almost 3 times greater than the defined threshold for functional ORFs.

**Fig. 4.**
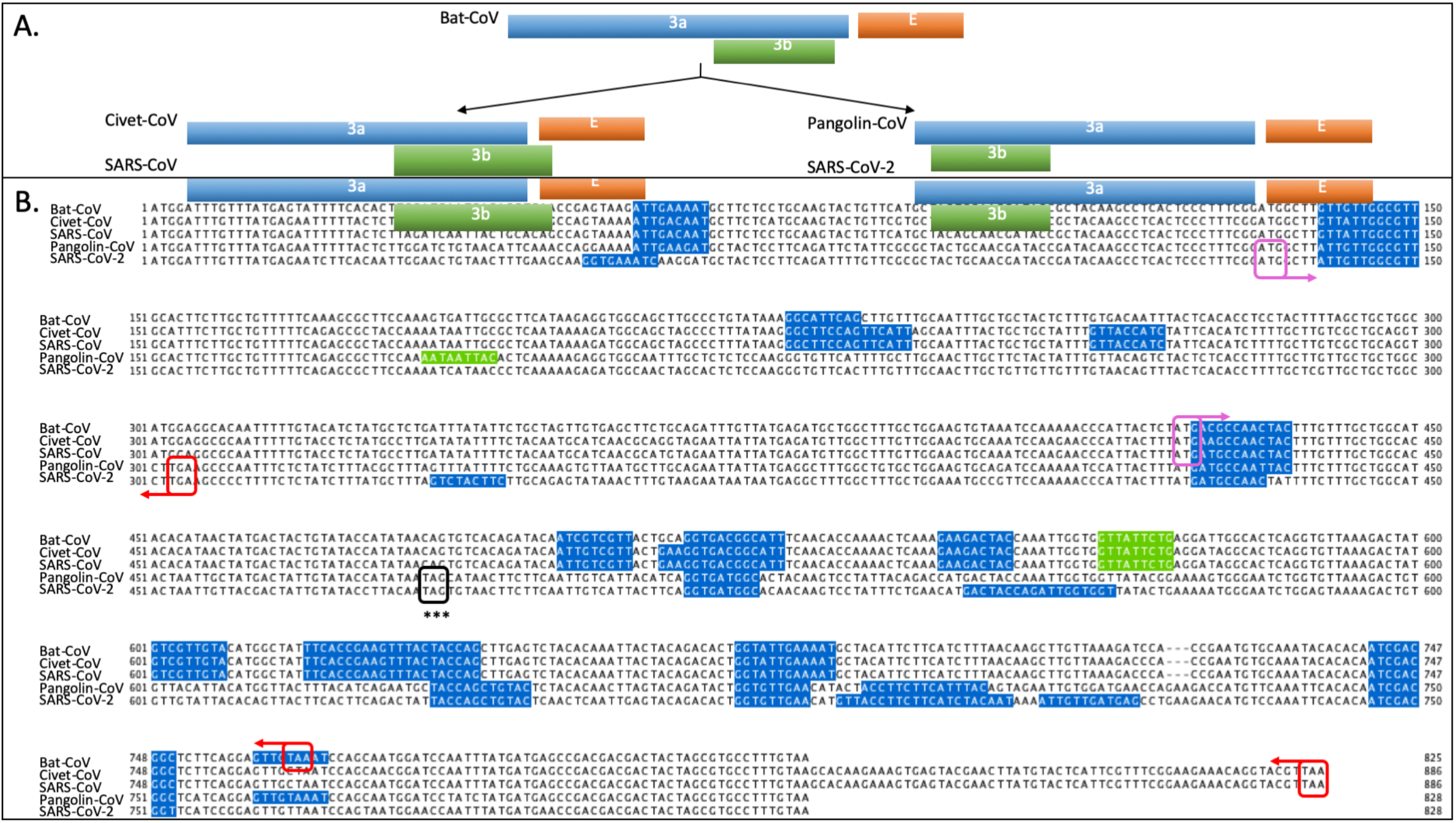
**A.** Schematic view of genome organization of ORF3a, ORF3b and E gene. **B.** Multiple alignment of ORF3a, ORF3b sequences, with *X* motifs in the reading frame of ORF3a shown in blue. The start and stop codons of the overlapping ORF3b sequences (in the +1 reading frame of ORF3a) are indicated by purple and red boxes respectively. *X* motifs in the reading frame of ORF3b are shown in green.

The ORF3b coding sequence overlaps the +1 reading frame of ORF3a and sometimes extends beyond the start codon of the E gene. In SARS-CoV, it is proposed to antagonize interferon (IFN) function by modulating the activity of IFN regulatory factor 3 (IRF3) (Kopecky-Bromberg et al., 2007). However, immunohistochemical analyses of tissue biopsies and/or autopsies of SARS-CoV-infected patients have failed to demonstrate the presence of ORF3b *in vivo*, and the presence of ORF3b in SARS-CoV-infected Vero E6 cells is the only evidence for the expression of this protein (McBride and Fielding, 2012). Furthermore, when mice are infected with mutant SARS-CoV lacking ORF3b, the deletion viruses grow to levels similar to those of wild-type virus, which demonstrates that SARS-CoV is able to inhibit the host IFN response without the 3b gene (Yount et al., 2005).

Bat-Cov and Civet-CoV also present ORF3b overlapping the 3’ region of ORF3a (start codon at nt 422), although the sequence of Bat-CoV ORF3b is shorter having a stop codon within the ORF3a sequence (nt 764). We observe a single *X* motif in the ORF3b reading frame of length 9 nucleotides (563-571), resulting in low XME scores of 2.6, 1.9 and 1.9 respectively for Bat-CoV, Civet-CoV and SARS-CoV ORF3b. This ORF is not predicted to be present in Pangolin-CoV or SARS-CoV-2 due to the introduction of a new stop codon (indicated by *** in Fig. 4) and the loss of the *X* motif in the +1 reading frame.

However, a completely different ORF is identified in the Pangolin-CoV and SARS-CoV-2 sequences, overlapping the 5’ region of ORF3a (132-305). This ORF is not annotated in the SARS-CoV-2 reference genome (MT072688), but is annotated as ORF3b in the genome of another SARS-CoV-2 strain isolated from the first U.S. case of COVID-19 (MN985325). The Pangolin-CoV ORF3b sequence contains one *X* motif in the reading frame of length 9 nucleotides (183-191), but the *X* motif is lost in the SARS-CoV-2 genome.

### ORF8: a rapidly evolving region of SARS-CoV genomes

Previously shown to be a recombination hotspot, ORF8 is one of the most rapidly evolving regions among SARS-CoV genomes (Ceraolo and Giorgi, 2020). Furthermore, the evolution of ORF8 is supposed to play a significant role in adaptation to the human host following interspecies transmission and virus replicative efficiency (Xu et al., 2020).

In SARS-CoV isolated from bats and civets (as well as early human isolates of the SARS-CoV outbreak in 2003: data not shown), ORF8 encodes a single protein of length 122 amino acids (Fig. 5). However, in SARS-CoV isolated from humans during the peak of the epidemic, there is a 29-nt deletion in the middle of ORF8, resulting in the splitting of ORF8 into two smaller ORFs, namely ORF8a and ORF8b (Oostra et al., 2007). ORF8a and ORF8b encode a 39 amino acid and 84 amino acid polypeptide, respectively. The XME scores in these ORFs are in line with the known experimental evidence concerning their functions. ORF8a has an XME score of 15.3 in SARS-CoV and anti-p8a antibodies were identified in some patients with SARS (Chen et al., 2007). In contrast, ORF8b has no *X* motifs in the reading frame, and protein 8b was not detected in SARS-CoV-infected Vero E6 cells (Oostra et al., 2007).

**Fig. 5.**
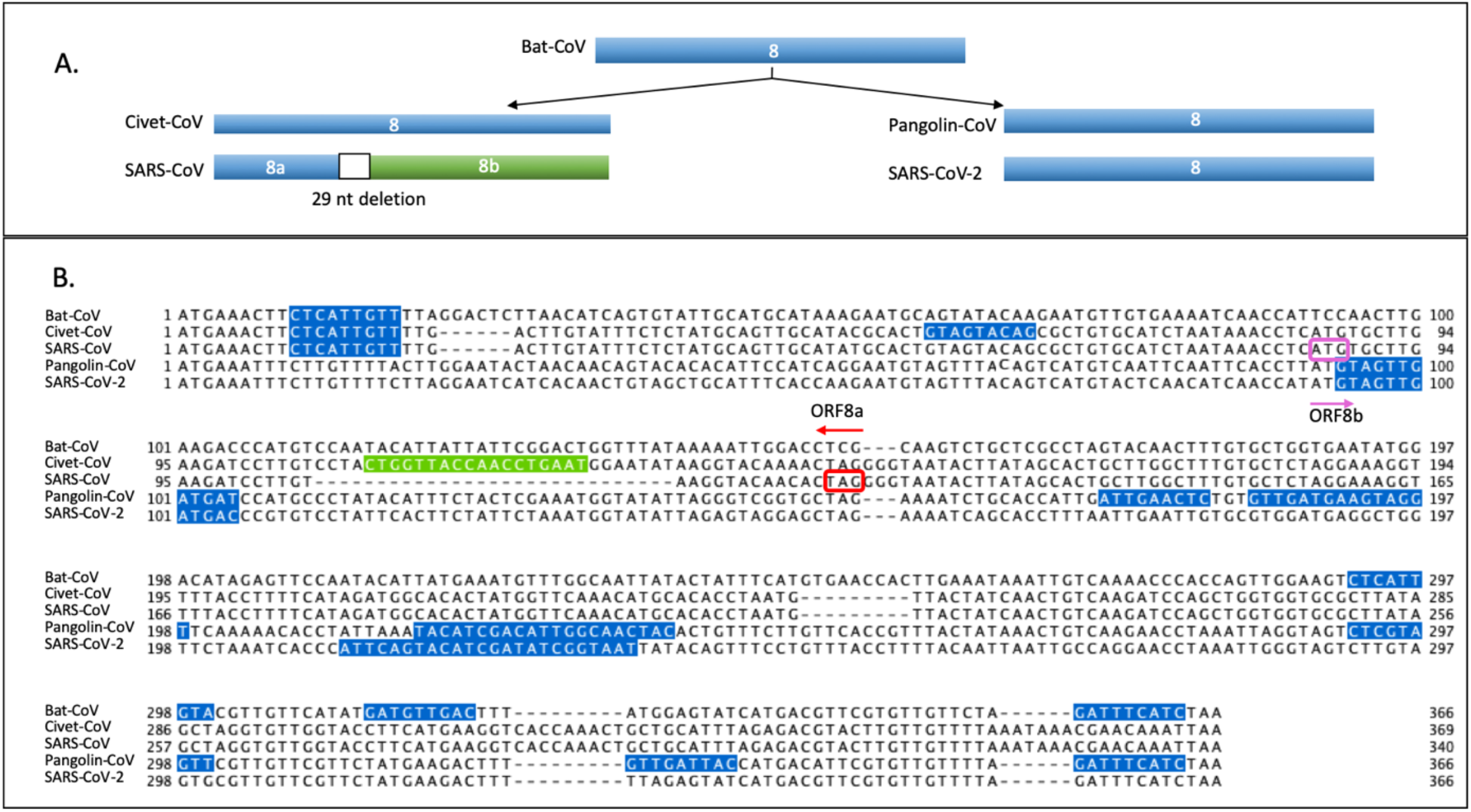
**A.** Schematic view of genome organization of ORF8, highlighting the 29-nt deletion in SARS-CoV, resulting in 2 ORFs: ORF8a and ORF8b. **B.** Multiple alignment of ORF8 sequences, with *X* motifs in the reading frame of ORF3a shown in blue. The start and stop codons of the SARS-CoV ORF8a and ORF8b sequences are indicated by purple and red boxes respectively. The *X* motif corresponding to the 29-nt deletion is shown in green.

It is interesting to note that although Civet-CoV has a full-length ORF8, it has a low XME score (XME=4.9) compared to Bat-CoV (XME=9.9). Thus, it is tempting to suggest that the loss of *X* motifs in transmission of the virus from bats to civets is somehow linked to the loss of ORF8 in the transmission from civets to humans. Both Pangolin-CoV and most SARS-CoV-2 strains contain the full length ORF8, with XME scores of 23.1 and 12.4 respectively. However, a 382-nt deletion has been reported recently covering almost the entire ORF8 of SARS-CoV-2 obtained from eight hospitalized patients in Singapore, that has been hypothesized to lead to an attenuated phenotype of SARS-CoV-2 (Su et al., 2020).

### Characterization of ORFs overlapping the N gene

The annotation of functional ORFs overlapping the N gene is variable in the different genomes studied here. In SARS-CoV, only ORF9b has been observed to be translated, probably *via* a ribosomal leaky scanning mechanism and may have a function during virus assembly (Xu et al., 2009; Shukla and Hilgenfeld, 2015). ORF9b limits host cell interferon responses by targeting the mitochondrial-associated adaptor molecule (MAVS) signalosome. However, some SARS-CoV strains have an additional ORF9c, annotated as a hypothetical protein (e.g. Genbank:AY274119). For Bat-CoV and Pangolin-CoV, no overlapping genes are annotated in the corresponding Genbank entries. In contrast, the Civet-CoV genome is predicted to contain both overlapping genes, ORF9b and ORF9c. Similarly, the annotation of overlapping ORFs for SARS-CoV-2 is different depending on the strain: the reference strain has no overlapping ORFs of the N gene, while the U.S. strain has ORF9b and ORF9c (see Methods). ORF9c is described as a short polypeptide (70 amino acids) dispensable for viral replication, but there is no data yet providing evidence that the protein is expressed during SARS-CoV-2 infection.

Here, we predict that ORF9b and ORF9c are present in all genomes as overlapping ORFs within the N gene (Fig. 6). Furthermore, Pangolin-CoV may also have an additional ORF, that we called ORF9d (XME=12.7), in the 3’ region of the N gene.

**Fig 6.**
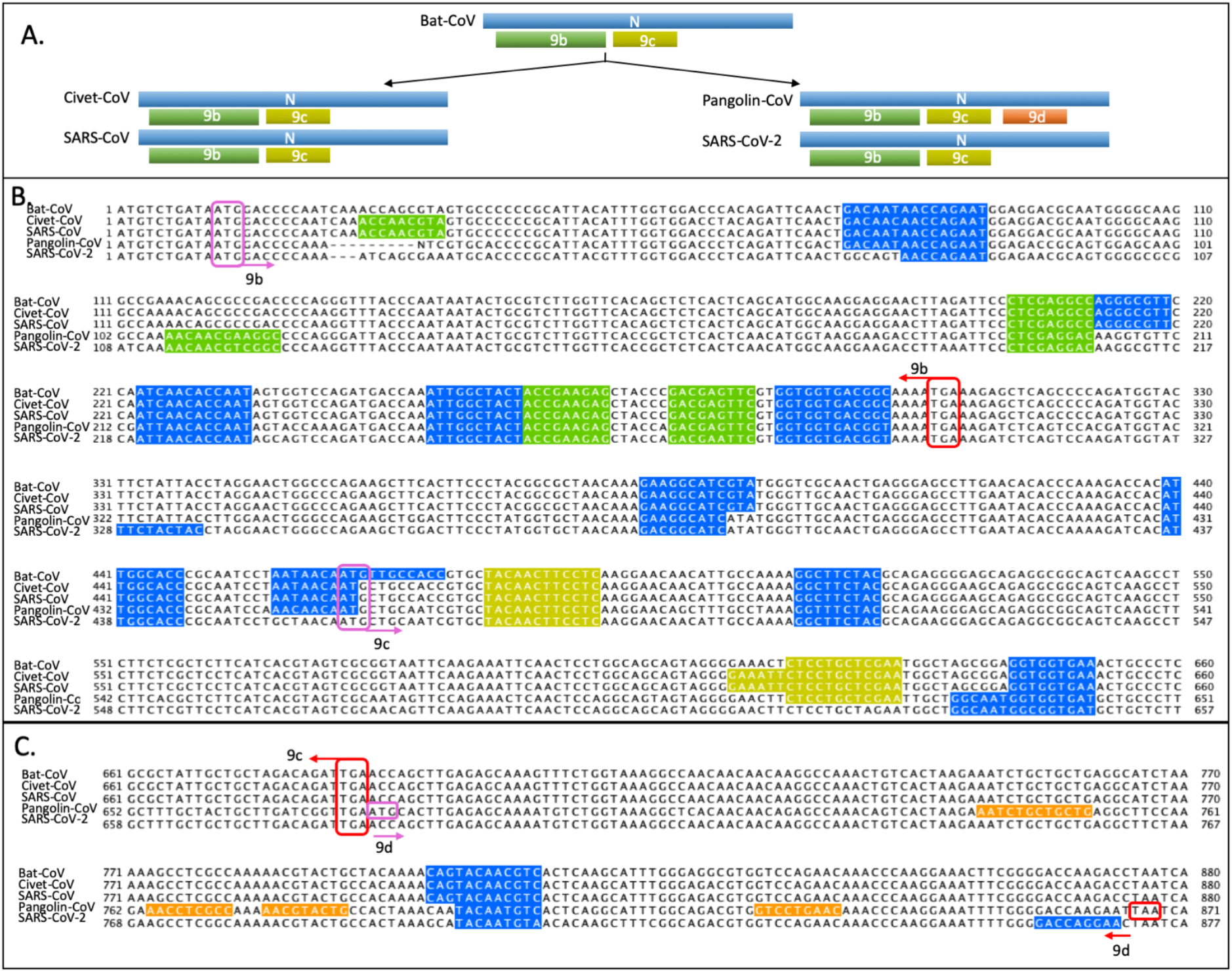
A. Schematic view of genome organization of ORF N, with overlapping genes ORF9b, 9c and the novel predicted 9d. B. Multiple alignment of ORF N sequences, with *X* motifs in the reading frame of ORF N shown in blue, in ORF9b in green, in ORF9c in yellow. Start and stop codons of the overlapping genes are indicated by violet and red boxes, respectively. C. The novel ORF9d predicted in Pangolin-Cov with *X* motifs in the reading frame shown in orange.

### Origin and evolution of ORF10

ORF10 is proposed as unique to SARS-CoV-2 (Wu et al., 2020) and codes for a peptide only 38 amino acids long. There is no data yet providing evidence that the protein is expressed during SARS-CoV-2 infection. Therefore, we wanted to investigate the potential origin of this protein. New proteins in viruses can originate from existing proteins acquired through horizontal gene transfer or through gene duplication for example, or can be generated *de novo*. To determine whether homologs of ORF10 are present in the other coronavirus genomes, we relaxed the GOFIX parameters used to predict functional ORFs, and set the minimum ORF length to 60 nucleotides. The predicted ORFs in the different genomes are shown in Fig. 7. The Pangolin-CoV genome contains a full-length ORF10 with XME=10.4, compared to the SARS-CoV-2 ORF10 with XME=20.2. A truncated version of ORF10 coding 26 amino acids is also detected in the Bat-CoV, Civet-CoV and SARS-CoV genomes, although this short ORF is probably not functional. We suggest that the ORF10 of SARS-CoV-2 thus evolved *via* the mutation of a stop codon (TAA) at nt 76 and the addition of a new *X* motif of length 15 nucleotides in the 3’ region.

**Fig 7.**
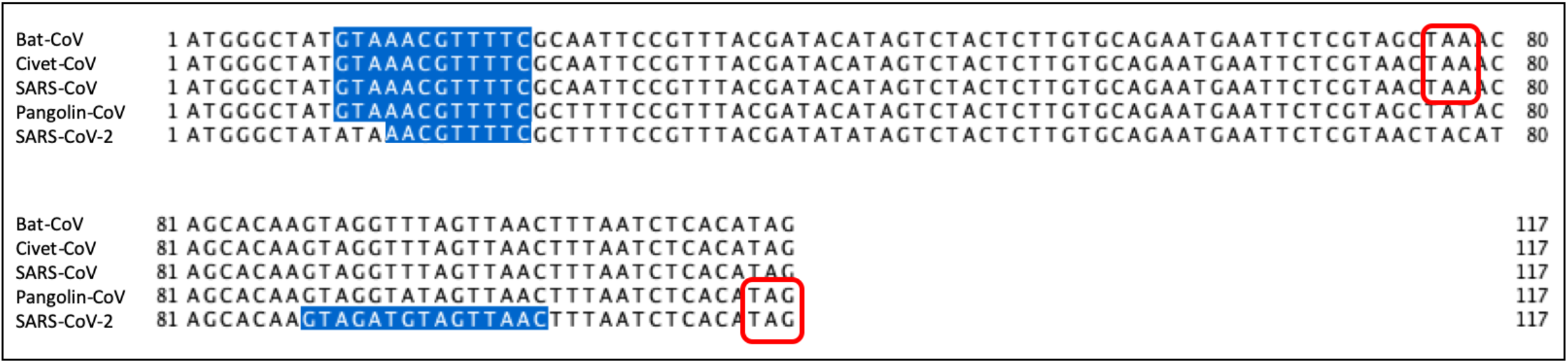
Multiple alignment of ORF10 sequences, with *X* motifs in the reading frame shown in blue. Stop codons are indicated by red boxes.

### Novel ORF overlapping the S gene

The GOFIX method predicts a novel ORF, that we called ORFSa, overlapping the RBD (Receptor Binding Domain) of the S (Spike) ORF in SARS-CoV (XME=7.6) and Civet-CoV (XME=7.6). ORFSa is found in the +1 frame and codes for a protein with 64 amino acids, as shown in Fig. 8. As the ORFSa sequence was not present in the Bat-CoV reference genome, we also searched for the ORF in the genomes of other Bat-CoV strains, and found one occurrence (XME=6.5) in the strain WIV16 (Genbank:KT444582) (Fig. 8), another bat coronavirus that is closely related to SARS-CoV (Yang et al., 2015).

**Fig. 8.**
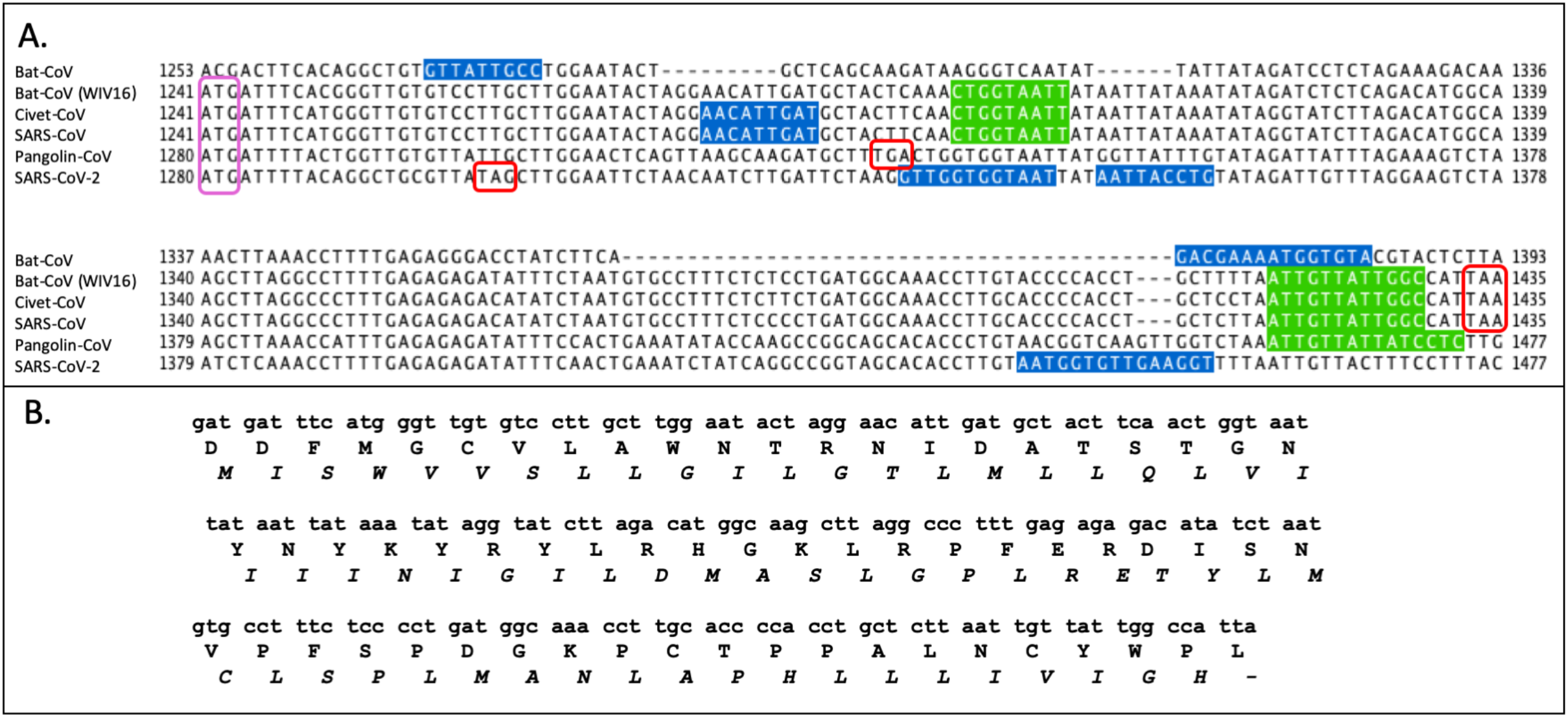
A. Multiple alignment of ORFSa sequences, with *X* motifs in the reading frame of ORF S shown in blue and ORFSa in green. Start and stop codons of the overlapping genes are indicated by violet and red boxes, respectively. Bat-CoV (WIV16) sequence is from Genbank:KT444582. B. Nucleotide and amino acid sequences of the novel ORF predicted to overlap the Spike protein in the genome of SARS-CoV. The nucleotide sequence segment (SARS-CoV:nt 22732-22926) encodes part (residues 414-478) of the RBD (residues 323-502) of the Spike protein (normal characters), while the reading frame +1 encodes a potential overlapping ORF (italics), which we named Sa.

To investigate whether the novel ORFSa might be a functional protein in SARS-CoV, we used BlastP to search the Genbank database for matches to viral proteins. A significant hit was obtained with a sequence identity of 100% to the protein AAR84376, described as “putative transmembrane protein 2d” from the genome of SARS coronavirus strain ZJ01 (AY28632). To further characterize this putative protein, the Phoebius web site (phobius.sbc.su.se) was used to predict transmembrane (TM) helices. Two potential TM helices of nearly twenty amino acids (residues 6-28 and 42-62) were predicted with a small inter-TM endodomain. Thus, this potential double-membrane spanning small protein might complement the set of already known SARS-CoV membrane proteins, namely the Spike (S), membrane (M) and envelope (E) proteins.

## Discussion

Coronaviruses are complex genomes with high plasticity in terms of gene content. This feature is thought to contribute to their ability to adapt to specific hosts and to facilitate host shifts (Cui et al., 2019). It is therefore essential to characterize the coding potential of coronavirus genomes. Here, we used an *ab initio* approach to identify potential functional ORFs in the genomes of a set of representative SARS or SARS-like coronaviruses. Our method allows comprehensive annotation of all ORFs. Surprisingly, the calculation of *X* motif enrichment is also accurate for the detection of overlapping genes, even though the codon usage and amino acid composition of overlapping genes is known to be significantly different from nonoverlapping genes (Pavesi et al., 2018).

We showed that the predictions made by the GOFIX method have high sensitivity and specificity compared to the known functional ORFs in the well characterized SARS-CoV. For example, the annotated ORFs that have been described previously as non-functional or redundant, notably ORF3b and ORF8b, are not predicted to be functional by GOFIX. In contrast, we identified a putative small ORF overlapping the RBD of the Spike protein in SARS-CoV, that is conserved in Civet-CoV and Bat-CoV strain WIV16. Protein sequence analysis predicts that this novel ORF codes for a double-membrane spanning protein.

We then used the method GOFIX to compare all putative ORFs in representative genomes, and showed that most are conserved in all genomes, including the structural proteins (S, E, M and N) and accessory proteins 3a, 6, 7a, 7b, 9b and 9c. However, a number of ORFs were predicted to be non-functional, notably ORF8b in SARS-CoV and ORF3b in all genomes. We also identified potential new ORFs, including ORF9d in Pangolin-CoV and ORF10 in all genomes.

Concerning SARS-CoV-2, to date, the coding potential of SARS-CoV-2 remains partially unknown, and distinct studies have provided different genome annotations (Zhou et al., 2020; Chan et al., 2020). Overall, the genome of SARS-CoV-2 has 89% nucleotide identity with bat SARS-like-CoV (ZXC21) and 82% with that of human SARS-CoV (Chan et al., 2020). Our analysis shows that the genome organization is conserved, and in particular ORF9b and ORF9c are predicted to be expressed in SARS-CoV-2 genome. As expected, the structural proteins, S, E, M and N are conserved and have similar XME scores. Here, we have shown that ORF3a, ORF6 and ORF9b in SARS-CoV-2 also have similar XME scores to SARS-CoV.

Previously identified differences include some interferon antagonists and inflammasome activators encoded by SARS-CoV that are not conserved in SARS-CoV-2, in particular ORF8 in SARS-CoV-2 and ORF8a,b in SARS-CoV, as well as the completely different ORF3b (Yuen et al., 2020). ORF3b has 0 *X* motifs in SARS-CoV-2 and expression was not observed in recent experiments aimed at characterizing the functions of SARS-CoV-2 proteins (Gordon et al., 2020). ORF10 is supposed to be unique to SARS-CoV-2, however it is also present in the Pangolin-CoV genome and its origin can be traced back to the Bat-CoV, where a truncated ORF of 26 amino acids, also present in the civet and human SARS-CoV genomes, can be found. Here, we observe that ORF7a, ORF7b and ORF9c have reduced XME scores in SARS-CoV-2. It remains to be seen whether these differences reflect functional divergences between SARS-CoV and SARS-CoV-2.

## Acknowledgements

The authors would like to thank the BiGEst Bioinformatics Platform for assistance. This work was supported by the Agence Nationale de la Recherche (BIPBIP: ANR-10-BINF-03-02; ReNaBi-IFB: ANR-11-INBS-0013, ELIXIR-EXCELERATE: GA-676559) and institute funds from the French Centre National de la Recherche Scientifique, and the University of Strasbourg.

